# Lights, Camera, Mirrors, Action! Toolbox for 3D Analysis of High-rate Maneuvers Using a Single Camera and Planar Mirrors

**DOI:** 10.1101/2023.03.14.532636

**Authors:** Wajahat Hussain, Mahum Naveed, Asad Khan, Taimoor Hasan Khan, Muhammad Latif Anjum, Shahzad Rasool, Adnan Maqsood

## Abstract

The advent of high-speed cameras being made available for commercial use has facilitated the discovery of intriguing animal behaviors. However, analyzing animal kinematics in three dimensions necessitates using several high-speed cameras, which are specialized equipment and come at a significant expense. In this study, we utilize a conventional methodology of capturing synchronized multiple views using a solitary camera and planar mirrors. We introduce a user-friendly software package that facilitates the calibration of this economical setup and enables the three-dimensional (3D) analysis of high-rate maneuvers. Within *∼* 15 minutes, a single high-speed camera is calibrated and can be used to acquire 3D data. We accompany the toolbox with a detailed user guide and video tutorial (https://tinyurl.com/yckm9y6b) for ease of use. We accurately reconstruct the wing beats of multiple flies (dragonfly, butterfly).

## INTRODUCTION

In 2014, about 11% of papers published in *J. Exp. Biol*. included video analysis (Theriault et al., 2014). Most of these works included two or more cameras. Using multiple cameras helps in measuring the three dimensional (3D) locations of points of interest in the scene (a.k.a 3D reconstruction). In addition to multiple views, another exciting dimension is the camera’s speed. Highspeed cameras, having frame rates ≫30 frames per sec (FPS), unveil findings that are invisible to the naked eye. Combining multiple high-speed views reveals, perhaps, the most exciting behaviors. Koehler et al. (2012) demonstrated that the wings of dragonflies are not rigid planes; instead, they display a cambered profile. However, the cost of high-speed cameras is prohibitively expensive, constraining their widespread adoption and usage. Phantom–a famous high-speed video camera brand by Vision Research–can be priced $100,000 and up (Moynihan, 2014). Furthermore, in addition to hardware setup for high-speed 3D data capture, commercial software, available to process synchronized streams of data, are expensive ($10,000 to $100,000) and can only operate with limited frame rates (< 200 FPS) (Jensen et al., 2020). We present a simple accessory that converts any high-speed camera into multiple high-speed virtual cameras (views), which can be used for 3D analysis. We also provide the accompanying software (calibration code) under the GNU license.

Recently, multiple attempts have been made to create an affordable alternative for capturing 3D data at high speed. Chalich et al. (2020) proposed a solution to set up a *∼* 2000 FPS camera at a tempting offer of 800 USD. Given such deals, deploying multiple cameras seems realistic. Unfortunately, this economic setup requires the advanced ability to design and implement embedded (FPGA) solutions. Zhang et al. (2015) proposed a simple accessory, consisting of controllable LEDs, which enables 3D reconstruction using a single camera, utilizing a well-known approach known as shape-from-shading. LEDs change the shading of a static 3D object in a synchronized manner. However, this approach can only reconstruct static 3D objects. Srinivasan et al. (2018) used a simple calibration pattern, along with a single camera, to estimate the 3D trajectory of the subject in controlled settings. This approach requires *apriori* information about the subject (wingspan) and can only estimate a single 3D point.

Some approaches based on structured light require only a single high-speed camera (Deetjen et al., 2017; Hyun et al., 2018; Li and Zhang, 2018). A projector emits structured patterns which are captured by a synchronized camera. Firstly, these customized solutions proposed in the literature are not readily available compared to simple high-speed cameras. Furthermore, they require accurate synchronization between the projector and the camera. The synchronization mechanism required for multiple high-speed sensors itself incurs substantial expenses. Currently, attempts are being made to devise cost-effective alternatives for synchronizing multiple sensors operating at high speeds (Laurijssen et al., 2018). Catadioptric systems provide an affordable substitute for capturing multiple views using a single camera. Such systems comprise a refractive lens, an integral part of the camera, in conjunction with reflecting mirrors (which form an additional setup) (Hecht amd Zajac, 1974). The imagery captured through reflection is commonly referred to as a virtual view.

A diverse array of catadioptric systems can only be limited by imagination. Numerous catadioptric systems have been developed, incorporating various types of mirrors, such as planar, parabolic, elliptic, and hyperbolic mirrors, among others (Gluckman and Nayar, 2001; Nene and Nayar, 1998). Catadioptric setup has also been proposed for structured light sensors (Lanman et al., 2009; Akay and Akgul, 2014).

The utilization of catadioptric systems provides several benefits, such as offering a broad field of view, synchronized views, low setup cost, and possessing similar intrinsic calibration parameters across different views (Gluckman and Nayar, 2001). However, many practical challenges hinder the widespread use of these low-cost catadioptric setups.

The primary obstacle is constructing a formal model of the catadioptric system that allows multiple views to estimate the 3D geometry of the captured imagery (Nene and Nayar, 1998). Firstly, setting up such systems requires solid computer vision knowledge. Secondly, although some of these systems are well understood in literature (Nene and Nayar, 1998; Gluckman and Nayar, 2001; Rodrigues et al., 2010; Ying et al. 2012), transforming this detailed mathematical literature into a user-friendly toolbox is not trivial. Hence more user-friendly toolboxes are required for such systems as is the case for simple (pinhole) cameras.

The impact of the non-user-friendly aspect of catadioptric systems, despite their low cost, can be assessed by the success of Kinect, a consumer depth camera introduced by Microsoft almost a decade ago. The research community later utilized the Kinect camera, which was initially developed for gaming consumers and accompanied by sparse documentation. Consequently, several 3D sensing applications were created using these relatively expensive depth cameras compared to simple webcams and catadioptric systems (Shotton et al., 2012; Newcombe et al., 2011).

A large amount of 3D data was generated using Kinect (Lai et al., 2011; Silberman et al., 2012). With the availability of large 3D datasets, machine learning methods emerged, which learned to predict depth using a single view from a normal camera (Bansal et al., 2016). Recently deep learning has been shown to estimate a detailed human pose using a single image (Fang et al., 2021). However, the frame rate of Kinect is 30 FPS, and its high frame rate alternatives (Deetjen et al., 2017; Hyun et al., 2018; Li and Zhang, 2018) are not commercially available. Can we achieve similar detailed 3D motion estimation with flying organisms?

To this end, we provide a user-friendly toolbox for catadioptric setup with a high-speed camera and planar mirrors. We show that it takes around 15 minutes to calibrate and recover the 3D motion of a flying bug. Our calibration process leverages the well-known camera calibration toolbox (Bouguet, 1999). Therefore, no additional training is required to deploy our solution. Our software is MATLAB based. We provide detailed documentation and video tutorials for smooth setup.

We used our approach to reconstruct the wings of a dragonfly and a butterfly. Results demonstrate that we achieve high accuracy in 3D analysis.

## MATERIALS AND METHODS

### Experimental Setup

We used a single high-speed camera (Phantom v2012, Vision Research, Inc., Wayne, NJ, USA) with a lens focusing on 18-140 mm. Monochrome images are captured at 12,000 FPS with 1280 × 800 resolution. We used a monochromatic light (MultiLED LT-V9-15, GS Vitec GmbH, Bad Soden-Salmünster, Germany) with 24 High-power LEDs, 84 Watt, 24 Volt (7,700 Lumen total). Finally, we used two ordinary mirrors to form the chamber’s walls.

### Quick Calibration Guide

We use two simple household mirrors to capture multiple and simultaneous views from a single high-speed camera (Fig. 1a).

**Fig. 1.**
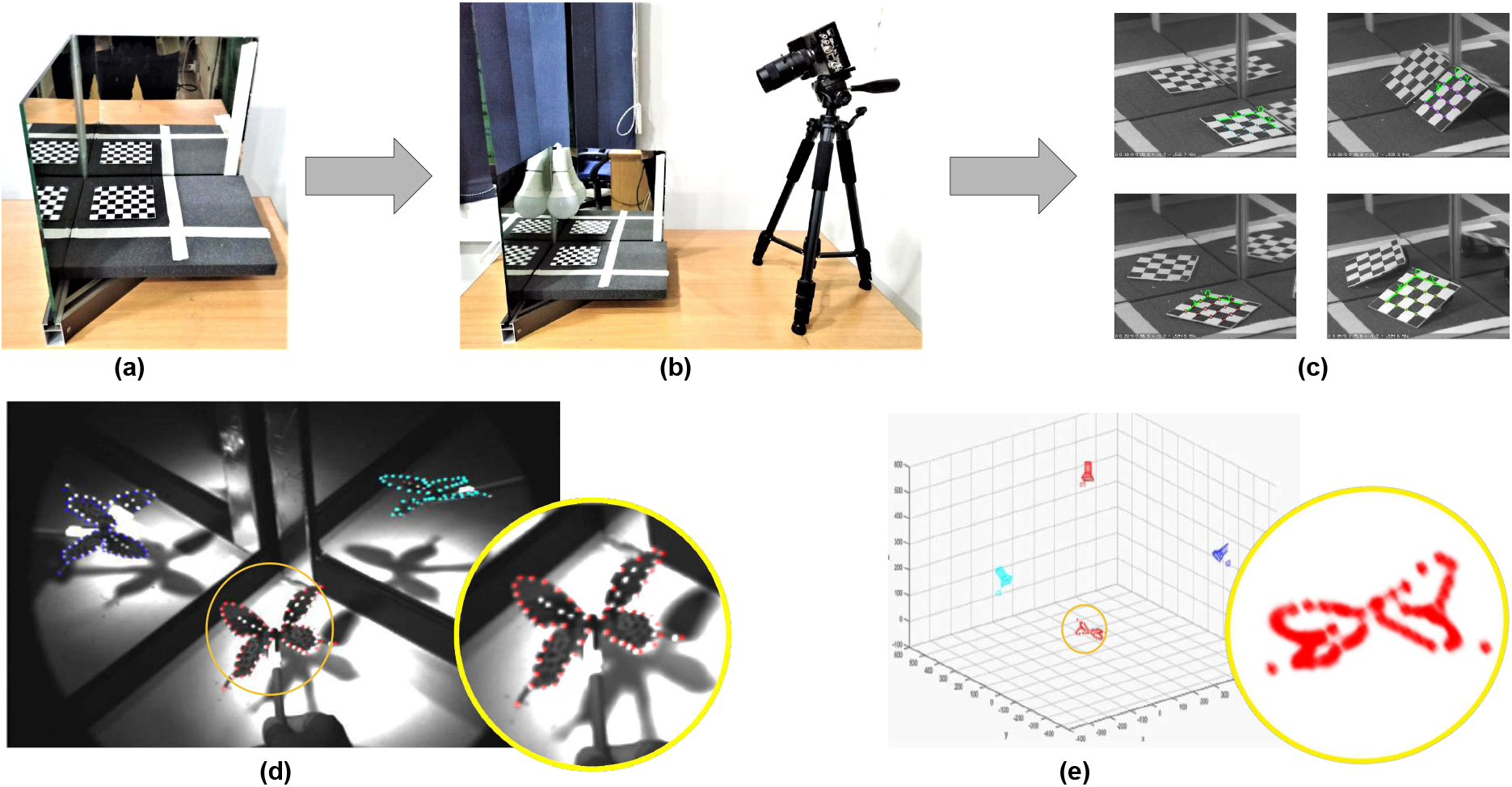
Calibration Setup: (a) Two household mirrors form the walls of the chamber to be used for housing the subject. A calibration pattern is placed on the floor of the chamber. (b) The camera is placed having the calibration pattern and its reflections in view. (c) The calibration pattern is moved to capture various shots. The rest of the setup remains static. (d) The target object and its reflections are captured in a single shot. (e) 3D reconstruction of the target object.

#### Algorithm 1

Calibration of Single Camera and Planar Mirrors

**Calibration Steps**:

1: Place two mirrors perpendicular to each other and the ground (Fig. 1a).

2: Place a checkered calibration pattern on the ground (Fig. 1a).

3: Place the camera on the tripod having the calibration pattern and its reflections in view (Fig. 1b).

4: Move the checker pattern to capture multiple (5-10) images (Fig. 1c).

5: Leave the camera and the mirror setup unmoved after capturing the last image. The checker pattern can be removed.

Complete the calibration using the guide provided on GitHub.

**Reconstruction Steps**:

6: Capture view of the target object similar to calibration pattern (Fig. 1d).

7: Reconstruct the target object using the toolbox (Fig. 1e).

Details are provided on the GitHub page.

These mirrors form the left and the center walls of the chamber housing the subject (Fig. 1a). Additional mirrors (ceiling, right side) can be added. In our experience, the perpendicular arrangement of mirrors gives the best calibration results (Fig. 2). This is convenient since the walls of the chamber are usually perpendicular. Transparent glass or plastic sheets may restrict the subject from escaping the chamber.

**Fig. 2.**
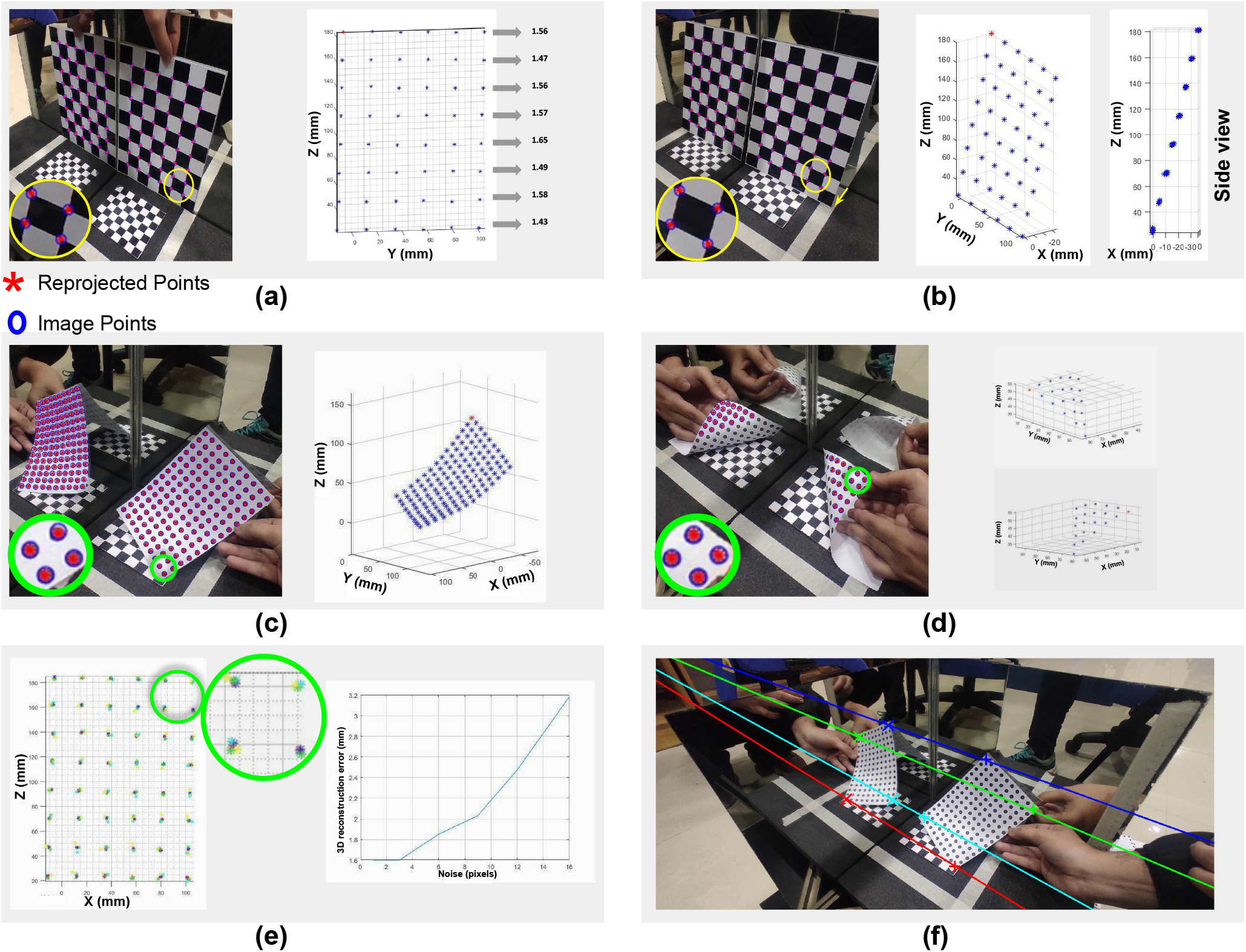
Accuracy Analysis. (a) **Vertical pattern:** checkered pattern, used for calibrating the setup, was mainly placed near the floor of the chamber. We reconstructed a vertical checker pattern to evaluate our approach’s accuracy for areas of the chamber not included in the calibration. We reconstructed 48 points with *∼* 1.50 mm RMS 3D reconstruction error and *∼* 2.12 pixels RMS reprojection error. (b) **Slanted pattern:** Similarly, we reconstructed the slanted checker pattern of arbitrary orientation and achieved *∼* 1.26 mm RMS 3D reconstruction error and *∼* 1.99 pixels RMS reprojection error. (c) **Concave pattern:** We reconstructed 148 points with *∼* 0.94 pixels RMS reprojection error. (d) **Convex pattern:** We reconstructed 21 points with *∼* 0.98 pixels RMS reprojection error. (e) **Landmark Localization Error:** Landmarks on animals generally span multiple pixels and, therefore, have inherent localization errors. To measure the effect of this ambiguity, we added different noise levels in the image points of the vertical checker pattern. We achieved *∼* 3.18 mm RMS 3D reconstruction error for the highest noise level. (f) **Epipolar Lines:** Our toolbox also provides epipolar lines (four colored lines) which pass through the corresponding points (four corners of the page). These lines help remove corresponding incorrect points which occur during automatic matching. The distance between the epipolar line and the corresponding points is *∼* 1 pixel RMS which indicates the accuracy of our calibration.

A single image captured by the camera contains multiple views (subject and its reflections) (Fig. 1d). All the views are captured in a single image. Therefore, this avoids the expensive synchronization step. However, this simultaneous capture results in limited resolution for each view (Fig. 1d). As shown in experiments, this setup allows enough resolution to capture the fine details, including the natural landmarks on butterfly wings (Fig. 3b), the angle between (leg) joints of housefly (Fig. 3c) and artificial landmarks on wings of dragonfly (Fig. 3a).

**Fig. 3.**
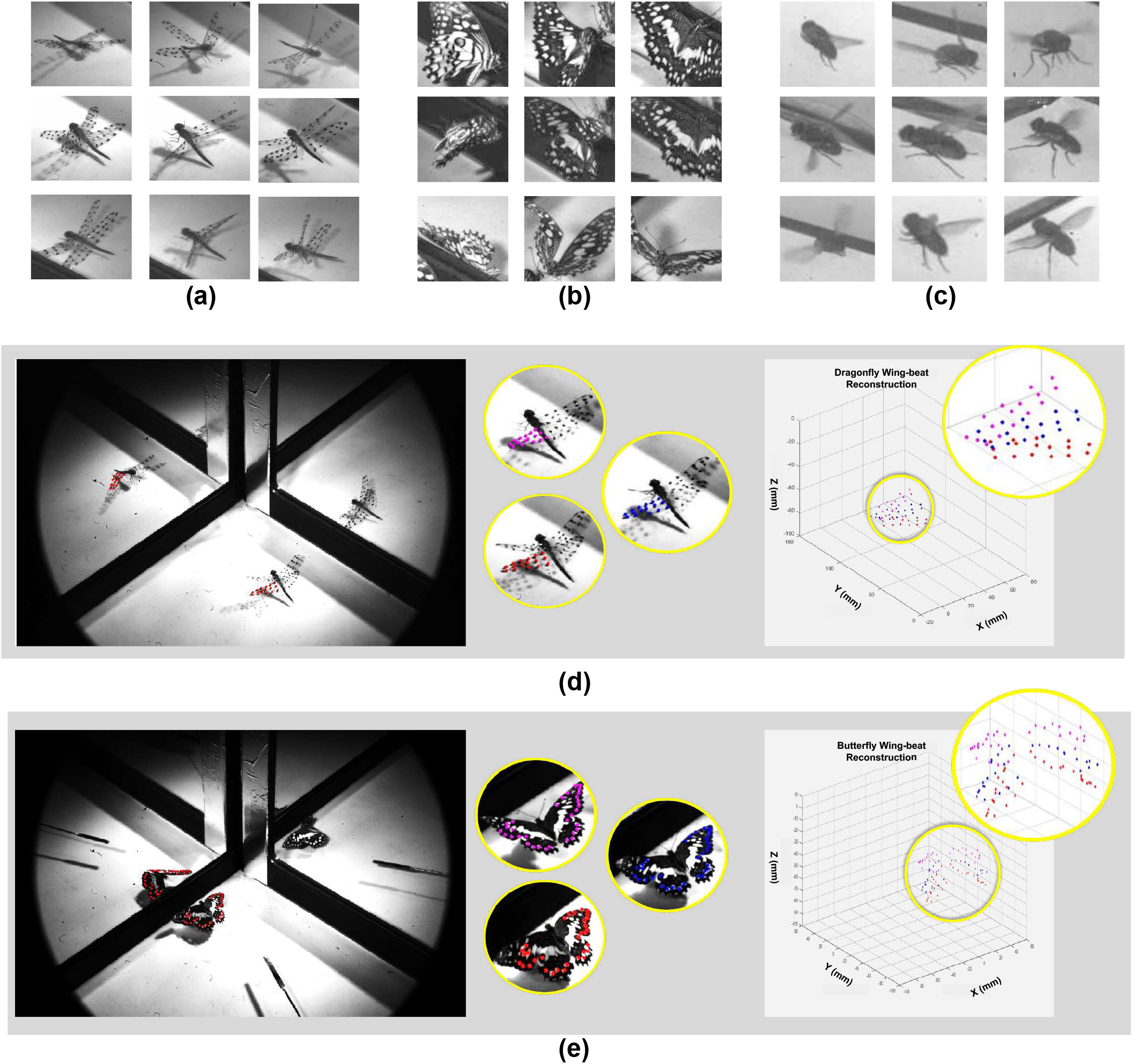
3D Reconstruction of Flies. Our proposed setup manages to capture multiple flies poses with fine details, including (a) artificial landmarks on dragonfly wings, (b) natural landmarks on butterfly wings, and (c) angle between leg joints of houseflies. (d) We reconstructed the wing beats of a dragonfly. The RMS reprojection error for 176 points was 1.98 pixels. Each spot was *∼* 16 pixels in diameter. The image size of the dragonfly’s wing was 169 pixels. (e) We also reconstructed the wing beats of a butterfly. The RMS reprojection error for 96 points was 4.72 pixels. Each spot was *∼*6.40 pixels in diameter. The image size of the butterfly’s wing was 46.5 pixels.

This initial setup and calibration require *∼* 15 minutes (Algorithm 1). We provide a user-friendly guide on setting up and using our toolbox on GitHub. A video tutorial accompanies it on YouTube.

Our calibration pattern generally moves along the ground plane (Fig. 1a). It is recommended to capture only a few views where the pattern is lifted from the ground plane (Fig. 1c). It is *not necessary* to move the calibration pattern in the entire space/volume of the chamber (Fig. 1a). As shown in experiments, we achieve 3D reconstruction accuracy of RMS *≤* 2 mm for objects far from the ground plane.

Another source of degradation is the localization of the landmark in the image. The landmarks (manually drawn spots on dragonflies (Koehler et al., 2012) and natural spots on butterflies) usually span multiple pixels. This makes it challenging to find accurate correspondences between multiple views of the same landmark. To evaluate this effect, we manually added noise of multiple levels (± 1 pixel up to ± 16 pixels), similar to Theriault et al., 2014, and provided 3D reconstruction accuracy analysis (Fig. 1a). We provide software to perform similar analyses for a larger volume. This will further assist users in planning their experiments, keeping in mind their accuracy demand.

Finally, automatic correspondence methods lead to data association errors. These correspondence errors can be removed using epipolar geometry (Hartely and Zisserman, 2003) constraints between real and virtual views. These constraints are also part of our provided toolbox (Fig. 1a).

## RESULTS AND DISCUSSION

The Institutional Review Board/Ethical Review Committee of the National University of Sciences and Technology, Pakistan (permit no. 03-2023-SEECS-01/1) approved the observation protocols utilized for examining flies. In this study, we present the results of our assessment of the accuracy of 3D reconstructions generated for multiple objects (i.e., planar, convex, and concave surfaces) and the wing beats of both dragonflies and butterflies.

We assessed the accuracy of our 3D reconstructions using two metrics. The first metric measured the difference (in millimeters) between the actual distance and the estimated distance between two 3D points. The second metric quantified the disparity (in pixels) between the image point and its re-projected point in the image. Both metrics were evaluated using the Root Mean Square (RMS) and reported in our results.

### 3D reconstruction using single camera

We calibrated the setup by moving the calibration pattern mainly on the ground plane (Fig. 1c). To evaluate the accuracy of our setup, for the part of the chamber not included during calibration, we reconstructed a vertical calibration pattern (Fig. 2a). We reconstructed 48 points with *∼* 1.50 mm RMS 3D reconstruction error and *∼* 2.12 pixels RMS reprojection error. Even for the first row of the calibration pattern, which is farthest from the ground plane, we achieve *∼* 2 mm RMS 3D reconstruction error. We also reconstructed the calibration pattern in an arbitrary orientation (Fig. 2b). We reconstructed 48 points with *∼*1.26 mm RMS 3D reconstruction error and *∼* 1.99 pixels RMS reprojection error.

We also reconstructed concave (Fig. 2c) and convex shapes (Fig. 2d). Wings of flies are not planar and exhibit similar twists. We reconstructed 148 points with *∼* 0.94 pixels RMS reprojection error for the concave shape. We reconstructed 21 points with *∼* 0.98 pixels RMS reprojection error for the convex shape.

To reconstruct a given image point, one has to select (click) the same entity in different views. Generally, natural (Fig. 3b) or artificial landmarks (Fig. 3c) span multiple pixels. This landmark localization ambiguity affects the reconstruction accuracy. We reconstructed a vertical calibration pattern by adding this noise in the selected image points (Fig. 3e). We added different noise levels (± 1 pixel up to ± 16 pixels). We achieved *∼* 3.18 mm RMS 3D reconstruction error for the highest noise level. This will further assist users in planning their experiments, keeping in mind their accuracy demand.

In this study, we performed a manual selection of corresponding points in different views, which can be a time-consuming process. An alternative approach is to use automatic correspondence methods, such as those proposed by (Lowe, 2004). However, automatic methods may produce data association errors that can lead to outlier points. To address this issue, we applied epipolar geometry constraints to remove any such outliers. (Fig. 2e). Epipolar lines accurately pass through four corners of the page in both the original and the reflected view. We achieve *∼* 1 pixel RMS distance between the corresponding points and the associated epipolar lines.

### 3D reconstruction of Dragonfly using single camera

To reconstruct the wing beats of the dragonfly, we employed our setup, as shown in (Fig. 3d). Our objective was to reconstruct the wings of the dragonfly with dense points. However, the reconstruction was challenging due to the lack of discriminant natural landmarks on the dragonfly’s wings. Like Koehler et al. (2012), we spotted the dragonfly’s wings (16 marks on each wing) using a black marker. The RMS reprojection error for 176 points was 1.98 pixels, and each artificial spot was approximately 16 pixels in diameter. The image size of the dragonfly’s wing was 169 pixels.

### 3D reconstruction of Butterfly using single camera

We also conducted a 3D reconstruction of the wing beats of a butterfly, as illustrated in (Fig. 3e). In this experiment, we reconstructed the distinctive features of the butterfly’s wings. The root mean square (RMS) reprojection error for 96 natural landmarks was *∼* 4.72 pixels. Each natural landmark was ~6.40 pixels in diameter. The size of the butterfly’s wing image was 46.5 pixels.

### CONCLUSION

Our innovative approach, referred to as the “single camera multiple views” solution, does not necessitate the purchase of additional costly hardware and instead utilizes a well-established standard 4 camera calibration method commonly used in the research community. To facilitate the adoption of this novel and cost-effective setup, we offer both the software and a video demonstration for calibration purposes. Our contribution is to enable researchers to fully exploit the potential of existing high speed cameras in their labs for high-rate 3D analysis.

## Acknowledgements

We are indebted to Dr. Majid Ali and Mr. Hassan Nazir (USPCAS-E department, NUST) for providing access to the high-speed camera and helping set up the experiments.

## Contribution

W. H.: developing methodology, writing the manuscript., M. N.: collecting data, developing methodology and software, A. K.: collecting data, evaluating methodology and software, T. K.: evaluating methodology and developing epipolar constraints, M.L.A.: evaluating toolbox and writing the manuscript, S. R.: study concept, writing manuscript. A. M.: study concept, assisting with hardware and related work.

## Data availability

Code, data, and video tutorials are available at https://tinyurl.com/yckm9y6b

## Supplementary

Insert the supplementary text here.

## REFERENCES

Theriault, D. H., Fuller, N. W., Jackson, B. E., Bluhm, E., Evangelista, D., Wu, Z., Betke, M. and Hedrick, T. L. (2014). A protocol and calibration method for accurate multi-camera field videography. J. Exp. Biol.. 217, 1843–1848.

Koehler, C., Liang, Z., Gaston, Z., Wan, H. and Dong, H. (2012). 3D reconstruction and analysis of wing deformation in free-flying dragonflies. J. Exp. Biol.. 215, 3018–3027.

Moynihan, T. (2014). A slo-mo camera with an insanely high frame rate (and price tag). Internet. Accessed July 2021, Available at https://www.wired.com/2014/07/phantom-v2511-camera/.

Chalich, Y., Mallick, A., Gupta, B. and Deen, M. J. (2020). Development of a low-cost, user-customizable, high-speed camera. PloS one. 15.

Zhang, Y., Gibson, G. M., Hay, R., Bowman, R. W., Padgett, M. J. and Edgar, M. P. (2015). A fast 3D reconstruction system with a low-cost camera accessory. Scientific reports. 5, 1–7.

Srinivasan, M. V., Vo, H. D. and Schiffner, I. (2018). 3D Reconstruction of Bird Flight Using a Single Video Camera. bioRxiv.

Deetjen, M. E., Biewener, A. A. and Lentink, D. (2017). High-speed surface reconstruction of a flying bird using structured light. J. Exp. Biol.. 220, 1956–1961.

Hyun, J.-S., Chiu, G. T.-C. and Zhang, S. (2018). High-speed and highaccuracy 3D surface measurement using a mechanical projector. Optics express. 26, 1474–1487.

Li, B. and Zhang, S. (2018). Novel method for measuring a dense 3d strain map of robotic flapping wings. Measurement Science and Technology. 29.

Laurijssen, D., Verreycken, E., Geipel, I., Daems, W., Peremans, H. and Steckel, J. (2018). Low-cost synchronization of high-speed audio and video recordings in bio-acoustic experiments. J. Exp. Biol.. 221.

Jensen, G. W., Van der Smagt, P., Heiss, E., Straka, H. and Kohl, T. (2020). SnakeStrike: A Low-Cost Open-Source High-Speed Multi-Camera Motion Capture System. Frontiers in Behavioral Neuroscience. 14.

Zhang, G., Sun, J., Chen, D., and Wang, Y. (2008). Flapping motion measurement of honeybee bilateral wings using four virtual structured-light sensors. Sensors and Actuators A: Physical. 148, 19–27.

Bouguet, J.-Y. (1999). Visual Methods for Three-Dimensional Modeling. PhD thesis, California Institute of Technology, Pasadena, CA, USA..

Balebail, S., Raja, S. K. and Sane, S. P. (2019). Landing maneuvers of houseflies on vertical and inverted surfaces. PloS one. 14, 3018–3027.

Nene, S. A. and Nayar, S. K. (1998). Stereo with mirrors. International Conference on Computer Vision. 1087–1094.

Gluckman, J. and Nayar, S. K. (2001). Catadioptric stereo using planar mirrors. International Journal of Computer Vision. 44, 65–79.

Rodrigues, R., Barreto, J. P. and Nunes, U. (2010). Camera pose estimation using images of planar mirror reflections. European Conference on Computer Vision. 382–395.

Ying, X., Peng, K., Hou, Y., Guan, S., Kong, J. and Zha, H. (2012). Self-calibration of catadioptric camera with two planar mirrors from silhouettes. IEEE transactions on pattern analysis and machine intelligence. 35, 1206–1220.

Akay, A. and Akgul, Y. S. (2014). 3D reconstruction with mirrors and RGB-D cameras. International Conference on Computer Vision Theory and Applications (VISAPP). 325–334.

Lanman, D., Crispell, D. and Taubin, G. (2009). Surround structured lighting: 3-D scanning with orthographic illumination. Computer Vision and Image Understanding. 113, 1107–1117.

Hecht, E. and Zajac, A. (1974). Optics addison-wesley. Reading, Mass. 19872, 350–351.

Fang, Q., Shuai, Q., Dong, J., Bao, H. and Zhou, X. (2021). Reconstructing 3d human pose by watching humans in the mirror. IEEE/CVF conference on computer vision and pattern recognition. 12814–12823.

Shotton, J., Girshick, R., Fitzgibbon, A., Sharp, T., Cook, M., Finocchio, M., Moore, R., Kohli, P., Criminisi, A., Kipman, A. and others (2012). Efficient human pose estimation from single depth images. IEEE transactions on pattern analysis and machine intelligence. 35, 2821–2840.

Newcombe, R. A., Lovegrove, S. J. and Davison, A. J. (2011). DTAM: Dense tracking and mapping in real-time. International conference on computer vision. 2320–2327.

Silberman, N., Hoiem, D., Kohli, P. and Fergus, R. (2012). Indoor segmentation and support inference from rgbd images. European conference on computer vision. 746–760.

Lai, K., Bo, L., Ren, X. and Fox, D. (2011). A large-scale hierarchical multiview rgb-d object dataset. IEEE international conference on robotics and automation. 1817–1824.

Bansal, A., Russell, B., and Gupta, A. (2016). Marr revisited: 2d-3d alignment via surface normal prediction. International conference on computer vision. 5965–5974.

Lowe, D. G. (2004). Distinctive image features from scale-invariant keypoints. International journal of computer vision. 60, 91–110.

Hartley, R. and Zisserman, A. (2003). Multiple view geometry in computer vision. Cambridge university press.

